# chopBAI: BAM index reduction solves I/O bottlenecks in the joint analysis of large sequencing cohorts

**DOI:** 10.1101/030825

**Authors:** Birte Kehr, Páll Melsted

## Abstract

**Summary:** Advances in sequencing capacity have lead to the generation of unprecedented amounts of genomic data. The processing of this data frequently leads to I/O bottlenecks, e. g. when analyzing a small genomic region across a large number of samples. The largest I/O burden is, however, often not imposed by the amount of data needed for the analysis but rather by index files that help retrieving this data. We have developed chopBAI, a program that can chop a BAM index (BAI) file into small pieces. The program outputs a list of BAI files each indexing a specified genomic interval. The output files are much smaller in size but maintain compatibility with existing software tools. We show how preprocessing BAI files with chopBAI can lead to a reduction of I/O by more than 95% during the analysis of 10 Kbp genomic regions, eventually enabling the joint analysis of more than 10,000 individuals.

**Availability and Implementation:** The software is implemented in C++, GPL licensed and available at http://github.com/DecodeGenetics/chopBAI

## 1 INTRODUCTION

Sequencing capacity has increased dramatically in recent years making it feasible to sequence cohorts of tens of thousands of individuals. The recent introduction of the HiSeq X Ten system allows for sequencing up to 18,000 whole human genomes per year at 30-fold coverage. Raw sequencing data is typically processed with standard bioinformatics pipelines on a computational cluster, parallelized by sample. Several variant calling tools, such as GATK’s UnifiedGenotyper (McKenna *et al.,* 2010), need to work with aligned reads from all sequenced individuals in order to achieve higher accuracy and comparability across samples. Instead of parallelizing by sample, we can parallelize this joint analysis by genomic region. Here, BAM index files allow quick random access to genomic regions, thus limiting the amount of I/O performed on BAM files that are typically more than 50 Gb in size.

Unfortunately this methodology does not scale for analyses of tens of thousands individuals. In many cases we are interested in small regions where the amount of data transferred per individuals is dwarfed by the size of the index file. A typical BAI index file is roughly 10 Mb in size, hence around 100 Gb of data transfer to a cluster node for each region is required in the case of 10,000 individuals. Depending on the size of the region this overhead can be an order of magnitude larger than the data transfer required to obtain the aligned reads from the BAM files. Thus, the transfer of the indices becomes an I/O bottleneck in the network.

To solve this issue we propose a method for chopping up the index in a predictable fashion, so that each cluster node can use a small portion of the overall index and network traffic is reduced significantly. We describe our method, chopBAI, and show a reduction in data transfer of more than 95 % for 10 Kbp genomic regions, while maintaining compatibility with existing software tools designed for indexed BAM files.

## 2 METHODS

Our program chopBAI implements a reduction of a BAM index file to a specified genomic interval. The resulting index is much smaller in size and is semantically equivalent to the complete index, in the sense that it will give the same answers for all queries to reads within the interval of interest; we make no guarantees to queries outside of the interval.

In the following, we first recapitulate the structure of a BAM index and how the reads are retrieved from a BAM file using a BAM index, before describing the reduction implemented in chopBAI and explaining the behaviour when using a reduced BAM index file.

### 2.1 BAM index structure.

BAM files store aligned reads in *chunks,* compressed sets of aligned reads where the size of the set is determined such that the uncompressed information fits into a predefined amount of memory. A BAM index makes use of the format of BAM files and allows for efficient navigation to chunks of aligned reads. As a result, we only need to decompress a small number of chunks and iterate through the aligned reads from only the beginning of chunks, instead of the beginning of the entire BAM file.

BAM index files (http://samtools.github.io/hts-specs/SAMv1.pdf) store file offsets of the beginnings and ends of chunks. The index files typically consist of three sections: metadata including a list of all reference sequences (chromosomes), bin indices (Kent *et al*., 2002) for all reference sequences, and linear indices for all reference sequences.

The *bin indices* are lists of bins, each bin storing the file offsets of a set of chunks. Bins represent contiguous genomic intervals that are of a predefined number of sizes. Any two bins do not partly overlap; either they are disjoint or one is completely contained in the other. Each chunk’s file offsets are stored in the smallest bin that fully contains all the alignments in the chunk. In addition to bin indices, BAI files contain *linear indices.* The linear indices store the smallest file offsets of aligned reads in all 16 Kbp windows tiling the reference sequences.

### 2.2 Retrieval of reads with a BAM index

The SAM specification provides C code for computing the list of bins in a bin index that overlap a genomic interval *I*. We refer to bins resulting from this computation as candidate bins. The set of chunks referenced in the candidate bins can further be filtered using a minimal file offset determined for *I* from the linear index. This is especially useful for the top level bins whose range spans 64 to 512 Mbp. Only the chunks of aligned reads remaining after this filtering need to be decompressed and further inspected to retrieve all reads whose alignments overlap *I*.

### 2.3 Reduction of a BAM index to an interval.

chopBAI achieves the reduction of an index to an interval *I* by considering which chunks would potentially be inspected during the retrieval of reads from *I* and any subset of *I*. Only these chunks are included in the reduced bin index. Following the algorithm for retrieving reads, chopBAI determines candidate bins and further narrows down the list of chunks within candidate bins using the information in the linear index. Optionally, chopBAI copies the linear index of the reference sequence of interest up to the end of *I*. It is important to note that this reduction only operates on the complete index itself and does not query the original BAM file.

### 2.4 Retrieval of reads with a reduced BAM index

The behaviour when querying a reduced BAM index file for a region that lies within the interval of the reduced BAM index does not differ from the behaviour on a full BAM index. The computation of bins that overlap the region returns only bins present in the reduced index; file offsets of all relevant chunks are present. In the case when the relevant part of the linear index has been copied, the set of chunks in the candidate bins can be filtered as usual. If the linear index was not copied in the reduction, it may be necessary to uncompress and iterate some additional chunks.

Querying a region outside the interval of the reduced BAM index does not result in an error. Instead, the region may appear as empty even though the BAM file contains aligned reads. It is left to the user to avoid such queries.

## 3 RESULTS

To evaluate the gain of preprocessing BAM indices with chopBAI we indexed a 62 Gb BAM file containing reads from an Icelander sequenced at 30-fold coverage on a HiSeqXTen sequencing machine and aligned with BWA-MEM (Li, 2013) to GRCh38. The complete BAM index is 8.8 Mb in size, whereas the reduced index built for a 1 Mbp region on chromosome 1 is on average only 4.5 Kb without linear index and 7.2 Kb including a linear index.

### 3.1 I/O reduction

To quantify chopBAI’s impact on I/O we measured the total amount of data transferred by `samtools` (Li *et al.,* 2009) when querying for regions of varying sizes. The data transfer was measured using the `strace` tool and includes reading the BAM index, BAM header and all chunks needed to retrieve the reads from the queried region.

Figure 1 displays the total amount of data read using the complete and reduced indices. The absolute difference in data transfer between the complete and reduced index remains the same over all tested interval sizes, but it becomes proportionally less with increasing interval size as shown by the log-scaled axis. We observe a 95 % reduction in the total data read for 10 Kbp regions, from 9.3 Mb for the full index to 450 Kb for the reduced index. The amount of data written does not change between the complete and reduced indices.

**Fig. 1.**
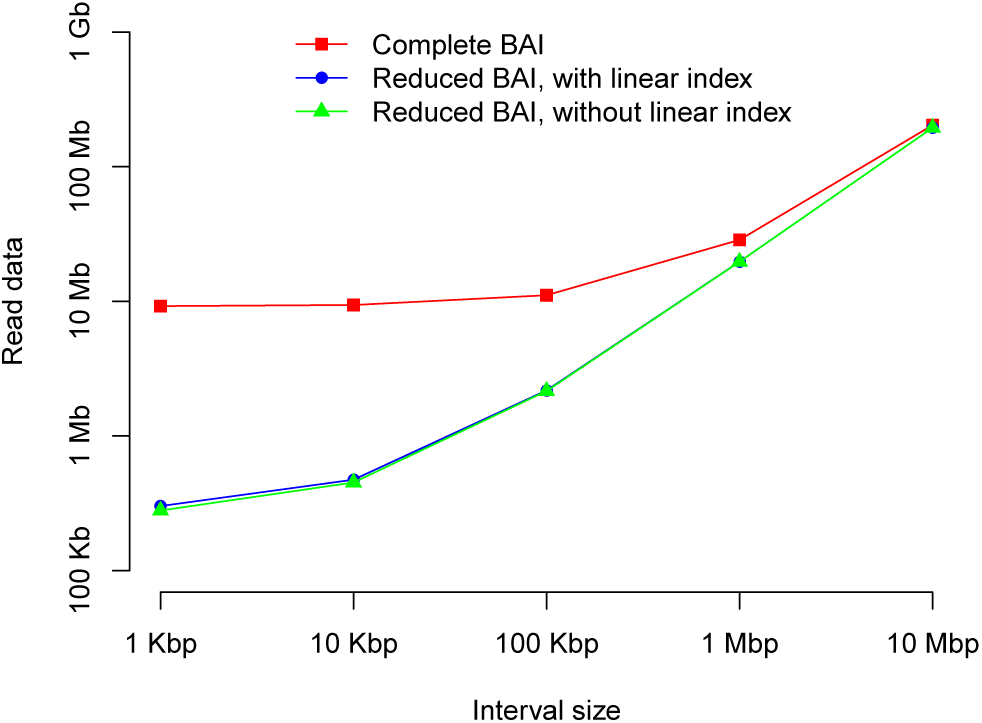
Amount of data transferred by `samtools` when querying genomic intervals of different sizes with the complete and with reduced BAI files. Indices were reduced to exactly the queried intervals. Averages over all intervals tiling chromosome 1 (10 Mbp and 1 Mbp) or all intervals tiling chr1:60,000,000-70,000,000 (100 Kbp, 10Kbp, 1 Kbp) are shown.

### 3.2 Running time

chopBAI can create reduced indices for a list of regions, allowing us to chop up the complete index into smaller pieces. The running time for chopping the BAM index for the complete human genome into 1 Mbp indices that overlap by 500 Kbp was approximately 15 seconds on a standard desktop computer, thus imposing a negligible overhead in terms of preprocessing.

## 4 DISCUSSION

When running an analysis of several small regions across thousands of individuals, such as certain commands in GATK (McKenna *et al.,* 2010) and PopIns (Kehr *et al.,* 2015), the status quo puts an unnecessary burden on the network of a computational cluster. With chopBAI’s preprocessing, the I/O imposed by BAM index files in the analysis of 10 Kbp regions of 10,000 BAM files can be reduced from 93 Gb to 4.5 Gb per job. This reduction in network I/O removes a significant bottleneck when processing a large set of individuals over small regions and enables running an order of magnitude more jobs simultaneously.

